# Predicting cell fate commitment of embryonic differentiation by single-cell graph entropy

**DOI:** 10.1101/2020.04.22.055244

**Authors:** Jiayuan Zhong, Chongyin Han, Xuhang Zhang, Pei Chen, Rui Liu

**Affiliations:** School of Mathematics, South China University of technology, Guangzhou, 510640, China; School of Biology and Biological Engineering, South China University of technology, Guangzhou, 510640, China; School of Computer Science and Engineering, South China University of technology, Guangzhou, 510640, China

**Keywords:** single-cell graph entropy (SGE), critical transition, embryonic differentiation, dark gene, cell fate commitment

## Abstract

Cell fate commitment occurs during early embryonic development, that is, the embryonic differentiation sometimes undergoes a critical phase transition or “tipping point” of cell fate commitment, at which there is a drastic or qualitative shift of the cell populations. In this study, we presented a novel computational approach, the single-cell graph entropy (SGE), to explore the gene-gene associations among cell populations based on single-cell RNA sequencing (scRNA-seq) data. Specifically, by transforming the sparse and fluctuating gene expression data to the stable local network entropy, the SGE score quantitatively characterizes the criticality of gene regulatory networks among cell populations, and thus can be employed to predict the tipping point of cell fate or lineage commitment at the single cell level. The proposed SGE method was applied to five scRNA-seq datasets. For all these datasets of embryonic differentiation, SGE effectively captures the signal of the impending cell fate transitions, which cannot be detected by gene expressions. Some “dark” genes that are non-differential but sensitive to SGE values were revealed. The successful identification of critical transition for all five datasets demonstrates the effectiveness of our method in analyzing scRNA-seq data from a network perspective, and the potential of SGE to track the dynamics of cell differentiation.

## 1. Introduction

Complex systems may switch abruptly to a contrasting state through a critical transition [1]. In recent years, detecting critical transitions for general systems, such as ecosystems systems [2-3], climates systems [4-5], financial systems [6,7], and epidemic model [8-9], has drawn more and more attentions. In biomedical fields, the rapid growth of single-cell datasets has shed new light on the complex mechanisms of cellular heterogeneity. In these single-cell experiments, the cell fate commitment represents a critical state transition or “tipping point” at which complex systems undergo a qualitative shift. Characterizing and predicting such critical transition is crucial for patient-specific disease modeling and drug testing [10]. Recent studies provided a plethora of statistical quantities such as variance, correlation coefficient, and coordination of gene expression, to detect a cell fate transition of embryonic differentiation [10,11]. However, these statistical quantities mainly focused on the analyses at the gene expression level, while single-cell RNA sequencing (scRNA-seq) may offer more information of an insight into the cell-specific network systems. In contrast to gene expression, cell-specific network is a stable form against the time and condition [12], and thus reliably characterize the biological processes such as cell fate commitment. Such a network system is viewed as a nonlinear dynamical system with interacted variables/biomolecules, whose dynamics can be roughly divided into three stages, the before-transition stage, the critical stage at which cell fate commitment occurs, and the after-transition stage [13,14]. However, to characterize the dynamics of biological system and predict the critical stage from single-cell dataset is challenging. Comparing with conventional bulk-cell information, single-cell analysis suffers from high dimensional, noisy, sparse and heterogeneous samples.

In this study, from cell-specific network viewpoint, we presented a computational method, the single-cell graph entropy (SGE), to detect the signal of a critical transition or cell fate commitment during the embryonic differentiation process, and identify key genes that play important roles in embryonic development. The utilization of SGE is based on rewiring the cell-specific networks with statistical dependency, calculating a network entropy score for each localized network, combining and analyzing the dynamical change of the local indices (Fig. 1). Such method can be viewed as data transformation from the “unstable” gene expression of single cells to the relatively “stable” SGE value of gene associations (Figs. 1A-1B). This SGE value can be analyzed by any traditional scRNA-seq algorithm for cell clustering, dimension reduction and pseudo trajectory analysis by simply replacing the original gene expressions with the SGE values. Notably, the SGE method has capabilities beyond traditional expression-based methods, that is, SGE aims at exploring the dynamically differential information at a single-cell level, and thus identifying a critical stage during the progression of a biological system (Fig. 1C). Specifically, we detect the signature of an imminent critical transition by a significant increase of the SGE value, which indeed reflects the dynamic change of cell heterogeneity and coordination of gene expression. The proposed approach has been applied to five scRNA-seq embryonic differentiation datasets, including mouse embryonic fibroblasts (MEF) to neurons, neural progenitor cells (NPCs) to neurons, human embryonic stem cells (hESCs) to definitive endoderm cells (DECs), mouse hepatoblasts cells (MHCs) to hepatocytes and cholangiocytes cells (HCCs), and mouse embryonic stem cells (mESCs) to mesoderm progenitors (MPs) from the NCBI GEO database. For these embryonic time-course differentiation datasets, the predicted cell fate transitions agree with the observation in original experiments. In these applications, from the dynamic perspective, it is also demonstrated that SGE has better performances than original gene expression in temporal clustering of cells, that is, the clustering analysis based on SGE score accurately distinguishes the cell heterogeneity over time while the gene expression fails. Based on the temporal clustering by SGE, the cell-lineage trajectories can be presented to further study the cell differentiation paths. Besides, in the analysis of these single-cell datasets, SGE uncovers a few “dark” genes, which are non-differential in gene expression but sensitive to SGE score and may play important roles in embryonic development (Fig. 1D). Therefore, the SGE method provides a new way to analyze the scRNA-seq data, and helps to track the dynamics of biological systems from the perspectives of network entropy. The successful application of SGE validated its effectiveness in single-cell analysis.

**Figure 1:**
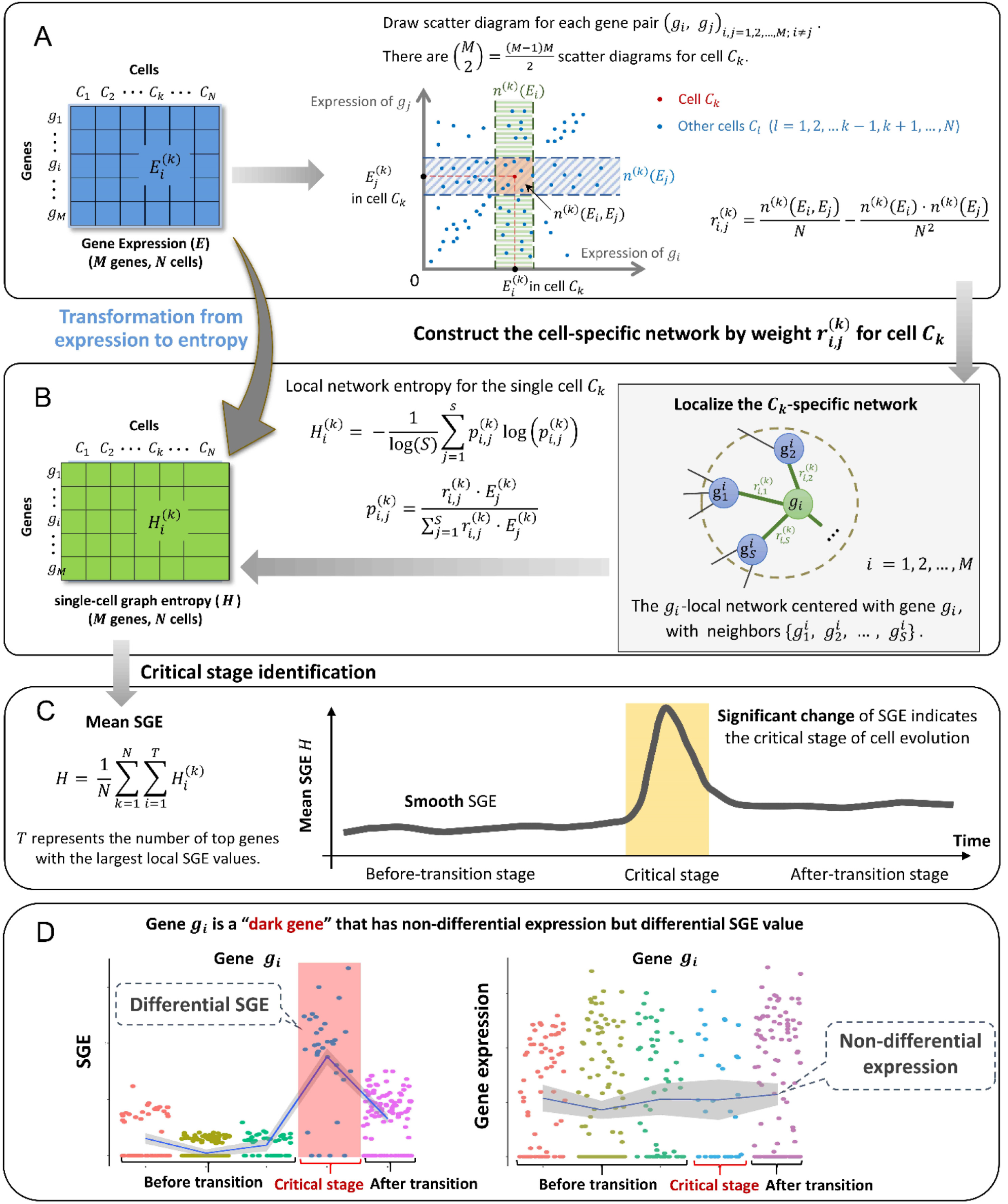
The schematic illustration of single-cell graph (SGE). **(A)** Draw scatter diagrams for every two genes, where each point represents a cell, and the expression values of the two genes in the *N* cells are mapped to the horizontal axis and the vertical axis respectively. Then *M* genes lead to *M* · (*M* − 1)/2 scatter diagrams. In the scatter diagram of genes *g*_*i*_ and *g*_*j*_, there is an edge between *g*_*i*_ and *g*_*j*_ in the cell *C*_*k*_ if the statistical dependency (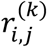 in Eq. (1)) is greater than zero, otherwise there is no edge. The *N*^(*k*)^(E_*i*_) and *N*^(*k*)^(E_*j*_) represent the number of the points (cells) in vertical box and horizontal box respectively. **(B)** Construct the cell-specific network by weight 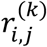 for cell *C*_*k*_. The*N* extract each local network from the cell-specific network. We calculate a local SGE score for each local network based on Eq. (2) and then get *M* local SGE scores corresponding to *M* local networks. **(C)** Critical transition can be predicted through the significant increase of SGE, i.e., the SGE keeps smooth when the system is in before-transition stage, while it increases abruptly when the system approaches the critical stage. **(D)** Different from the traditional biomarkers based on differential-expression genes, our SGE method uncovers some “dark genes”, which are sensitive to network entropy (SGE), but non-differential at the gene expression level.

## 2. Materials and Methods

### 2.1 Theoretical basis

A cell fate transition (cell fate commitment) occurs during the dynamical process of the early embryonic differentiation [10, 15-17]. Generally, the dynamical process of early embryonic development can be regarded as the evolution of a nonlinear dynamical system, while the cell fate transition is viewed as a drastic or qualitative state shift at a bifurcation point [10]. Similar to disease progression [13, 18], this dynamical process is modeled as three states or stages (Figure 1C): (1) a before-transition stage with high resilience; (2) a critical stage, which is the tipping point or cell fate transition with low resilience; (3) an after-transition stage, which is another stable state with high resilience.

In this study, the cell-specific networks were constructed based on a recently proposed statistical model [12], which provides a statistical dependency index (defined as Eq. (1)) to determine the gene associations at a single-cell level in a reliable manner. The statistic index ranges between −1 and 1. The positive statistical dependency value infers the statistically interacting relation between two genes, i.e., there is an edge between such two genes in the cell-specific network.

### 2.2 Algorithm to predict the critical transition based on SGE

Given the time series of single-cell RNA sequencing (scRNA-seq) data, the following algorithm is carried out to predict the critical transition.

**[Step 1]** At each time point, the logarithm log(1 + *x*) is applied to normalize the initial gene expression matrix with *M* rows/genes and *N* columns/cells, which is generated from the scRNA-seq data.

**[Step 2]** Constructing a specific network for each cell. Make scatter diagrams for every two genes in a cartesian coordinate system where the vertical- and horizontal-axes are the expression values of the two genes, respectively. For example, there are *N* plots in the scatter diagram for a gene pair (*g*_*i*_, *g*_*j*_) corresponding to the *N* cells. Each plot represents a cell, whose horizontal coordinate is 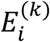 (the gene expression of *g*_*i*_ in cell *C*_*k*_) and the vertical coordinate is 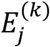(the gene expression of *g*_*j*_ in cell *C*_*k*_) (Fig. 1A). Then totally *M* · (*M* − 1)/2 scatter diagrams are obtained by making scatter diagram for every two genes. In the scatter diagram of genes *g*_*i*_ and *g*_*j*_, for the cell *C*_*k*_, whether there is an edge between *g*_*i*_ and *g*_*j*_ in the cell *C*_*k*_ is determined by the statistical dependency index as follows.

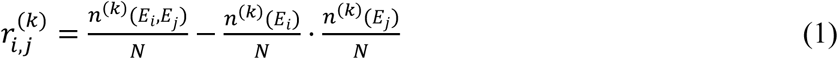

Two boxes near 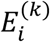 and 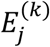 are drawn based on the predetermined integers such as 0.1*N*, which is proportional to the cell size *N*. The *N*^(*k*)^(*E*_*i*_) and *N*^(*k*)^(*E*_*j*_) represent the number of the points (cells) in vertical box, horizontal box respectively (Fig. 1A). We then straightforwardly obtain the third box, which is the overlapping of the previous two boxes. Therefore, the value of *N*^(*k*)^(*E*_*i*_, *E*_*j*_) can be obtained by counting the points (cells) in the third box. If the statistical dependency index i.e., Eq. (1) is greater than zero, there is an edge between *g*_*i*_ and *g*_*j*_ in the cell *C*_*k*_, otherwise there is no edge. By this way, we construct a cell-specific network *N*^(*k*)^ for cell *C*_*k*_, where each edge between two genes *g*_*i*_ and *g*_*j*_ is decided by the dependency index 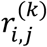.

**[Step 3]** Extracting each local network from the specific network. Specifically, for the cell *C*_*k*_, its specific network *N*^(*k*)^ can be segmented into totally *M* local networks. The local network 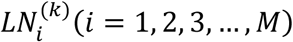 is centered at a gene *g*_*i*_, whose 1^st^-order neighbors 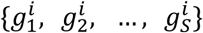 are the edges (Fig. 1B).

**[Step 4]** Calculating the local SGE value 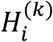 for each local network. Given the local network 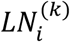 centered at a gene *g*_*i*_, its local SGE can be obtained as follow.

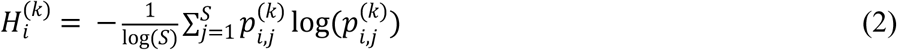

with

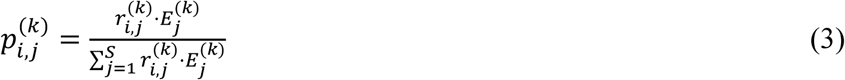

where 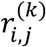 represents the weight coefficient between the center gene *g*_*i*_ and a neighbor 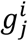, which is determined by Eq. (1). The value 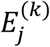 represents the gene expression of a neighbor 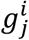 in *C*_*k*_ and constant *S* is the number of neighbors in the local network 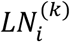. Clearly, the local SGE value (Eq. (2)) has been normalized to the number of nodes in a local network. After this step, the sparse gene expression matrix from the scRNA-seq data is transformed into a non-sparse graph entropy matrix (Figs. 1A and 1B), by taking the gene association into consideration. Thus, the local SGE value Eq. (2) is dependent not only on the expression of the center gene of a local network but also on the contribution from the neighboring genes.

**[Step 5]** Calculating the cell-specific SGE value *H*^(*K*)^ based on a group of genes with highest local SGE values, i.e.,

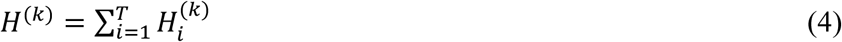

where constant *T* is an adjustable parameter representing the number of top 5% genes centered in its local networks with the highest local SGE values. In Eq. (4), *H*^(*k*)^ can be considered as the SGE score of the cell *C*_*k*_ and detect the early-warning signals of the cell fate transition. At each time point, the mean SGE score of a certain cell population is also employed in the tipping point detection. The mean SGE score of the top 5% genes with the largest local SGE values (Eq. (4)) was taken as the cell-specific graph entropy at a time point. In Supplementary_material_1 (Figure S1), it shows that different choices of *T* do not alter the identification of tipping point.

When the system approaches the vicinity of the critical point, the signaling genes or dynamical network biomarker (DNB) molecules exhibit obviously collective behaviors with fluctuations, which leads to that the dependent relations of DNB members in a critical transition state are different from those in a before-transition state. Moreover, the local SGE score 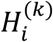 in Eq. (2) or the index *H*^(*K*)^ in Eq. (4) sharply increases when the system is near the critical stage (Fig. 1C). Thus, the SGE score can provide the early-warning signals of the cell fate transition.

## 3. Results

### 3.1 SGE predicting cell fate transitions for embryonic time-course differentiation

To demonstrate the effectiveness of SGE, the proposed method has been applied to five time-course datasets of embryonic differentiation from GEO database (http://www.ncbi.nlm.nih.gov/geo/), including MEF-to-Neurons data (ID: GSE67310) [19], NPCs-to-Neurons data (ID: GSE102066) [20], hESCs-to-DECs data (ID: GSE75748) [21], MHCs-to-HCCs data (ID: GSE90047) [22], and mESCs-to-MPs data (ID: GSE79578) [23]. The detailed description and sources of the datasets is given in Supplementary_material_1. The SGE score of each single cell was calculated according to the algorithm in Materials and Methods section. At each time point, the mean SGE score was taken to quantitatively measure the criticality of the cell population at this time point. An SGE curve across all time points was then employed to predict any possible cell fate transition of embryonic time-course differentiation.

For MEF-to-Neurons data, the mean SGE score abruptly increases from day 5 to day 20, as shown as the red curve in Fig. 2A. This significant change of SGE score provides the early-warning signal to an upcoming cell fate transition after day 20. This computational result agrees with the observation in original experiment, i.e., the differentiation of mouse embryonic intermediate cells into induced neuron (iN) occurs at day 22 [19]. Besides, to demonstrate the robustness of the proposed method in terms of the cells, the box plot of the cell-specific graph entropy was shown based on the samples of each time point. It is seen that the median values of the red box plot of SGE score in Fig. 2A also illustrates clear signal for the tipping point, which demonstrates that the SGE score is featured with robustness against sample noises. It is seen as the blue curve in Fig. 2A, the mean gene expression of the differential genes fails to provide any effective signals for cell fate transition. Therefore, the signature of a critical transition from MEF to neurons is identified by SGE at single-cell resolution of the cell populations.

**Figure 2:**
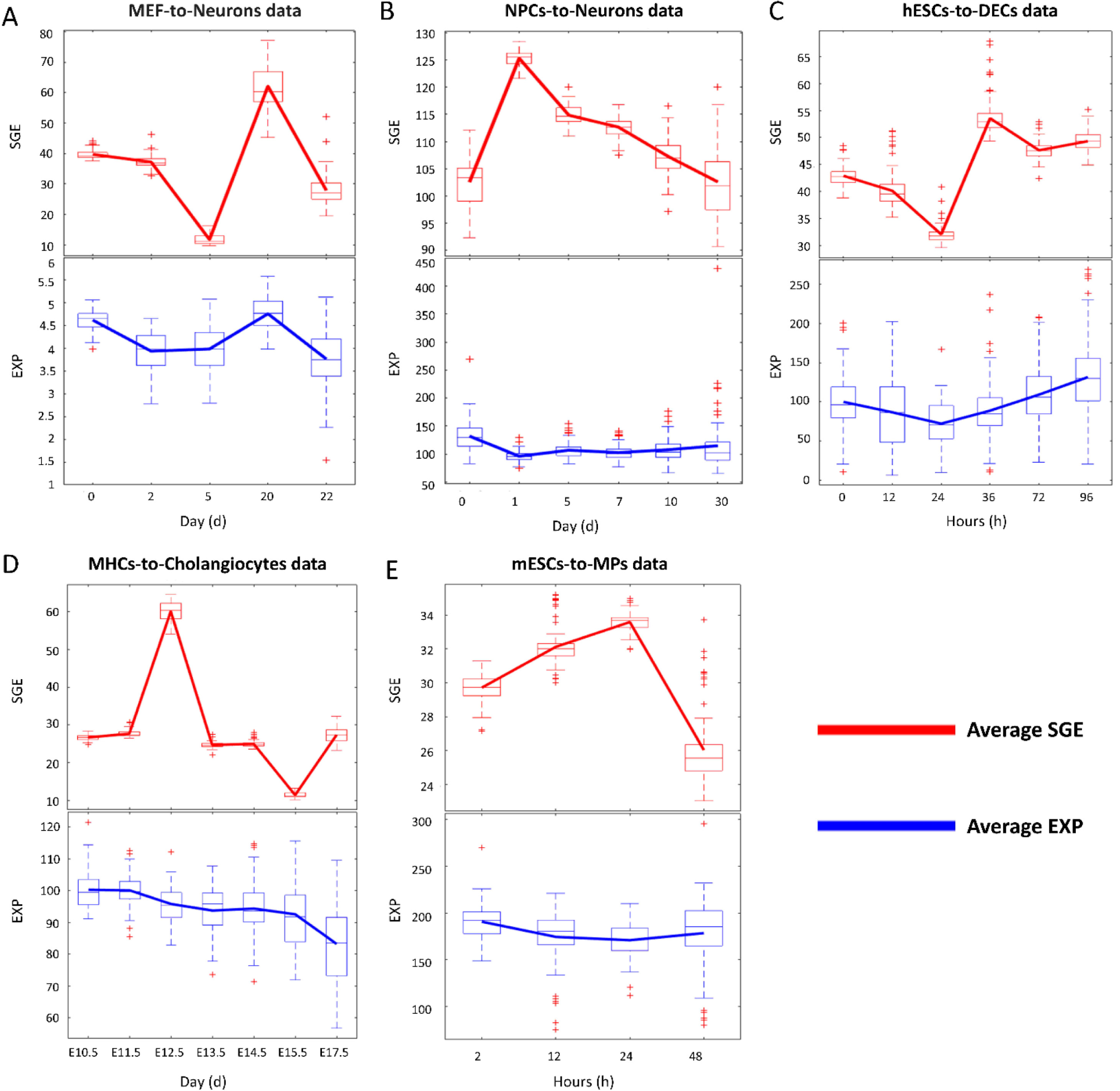
Predicting cell fate transitions in five embryonic differentiation datasets. **(A)** MEF-to-Neurons data **(B)** NPCs-to-Neurons data **(C)** hESCs-to-DECs data **(D)** MHCs-to-HCCs data and **(E)** mESCs-to-MPs data. The significant increase of SGE score as shown in the red curve indicates the imminent cell fate transition, while signaling genes at the gene expression level fails to provide any effective signals for the tipping point (the blue curve).

When applied to NPCs-to-Neurons data, i.e., a 30-day time-course differentiation experiment of neural progenitor cells into neurons, as shown as the red curve in Fig. 2B, the mean SGE score abruptly increases and reaches a peak at day 1, suggesting there is a cell fate transition after day 1. This signal also coincides with the observation in original experiment, in which it showed that the cells at day 1 were the least heterogeneous and after day 1 the transcriptional heterogeneity increased, reaching the largest heterogeneity among the neurons at day 30 eventually [20]. In addition, the median values of the red box plot of SGE score in Fig. 2B also demonstrated the robust performance of SGE score in detecting the early warning signal of a qualitative state transition. In contrast to SGE score, the mean gene expression fails to detect the early-warning signals of cell fate transition (the blue curve in Fig. 2B).

For hESCs-to-DECs data, the peak of the SGE score (the red curve in Fig 2C) appears at 36 h, which indicates an imminent cell fate transition after 36 h. Indeed, the differentiation induction into definite endoderm (DE) at 72 h, and the differentiation trajectory toward a DE fate commitment after 36 h, have been recorded in literatures [21,24], which validated the SGE signals. The robustness of SGE score in predicting the critical transition of the differentiation trajectory toward a DE fate can be showed by the median values of the box plot (the red box plot in Fig. 2C). Moreover, in terms of mean gene expression, there is no significant difference among six points time (the blue curve in Fig. 2C).

As the red curve shown in Fig. 2D, for MHCs-to-HCCs data, the drastic increase of average SGE appeared from E11.5 to E12.5 and reaches its peak at E12.5, after which hepatoblast-to-hepatocyte and cholangiocytes transition occurs [22]. Moreover, the median values of the red box plot of SGE score in Fig. 2D stably exhibits an obvious signal at the tipping point (E12.5), which demonstrates that SGE accurately predicts the cell fate transition for embryonic time-course differentiation. It is seen from the blue curve in Fig. 2D that the mean gene expression fails to provide any signal for the tipping point.

The SGE method has been applied to mESCs-to-MPs data, which is obtained from an experiment of a retinoic acid (RA)-driven differentiation of pluripotent mouse embryonic stem cells (mESCs) to lineage commitment [23]. It is seen from the red curve in Fig. 2E, the mean SGE score reaches its peak at 24 h, signaling an upcoming critical transition after 24 h. Actually, there are cells exiting from pluripotency between 24 h and 48 h of retinoic acid exposure and then differentiating into endoderm around 48 h [23]. Further, the median values of the box red plot of SGE score in Fig. 2E also indicates that the 24 h is a tipping point. But in terms of gene expression, it shows little significant difference among four points time (the blue curve in Fig. 2E).

The successfully prediction of the cell fate transitions during embryonic cell differentiation in these five datasets validates the effectiveness and accuracy of SGE method.

### 3.2 The dynamical change of local SGE scores

At the identified transition point, the group of top 5% genes with the largest local SGE values were taken as the signaling genes for further functional and biological analysis. These signaling genes can be regarded as a set of DNB and may be highly associated with the cell fate commitment during the embryonic development. First, the signaling genes were mapped to protein-protein interaction (PPI) network, from which the maximal connected subgraph was taken to study the dynamical network evolution. For MEF-to-Neurons data, we show the dynamical evolution of signaling genes at the network level (Fig. 3A). It is seen that a significant change of the network structure occurs at day 20, signaling an upcoming cell fate transition. Besides, the landscape of the local SGE score for signaling and non-signaling genes was illustrated as in Fig. 3D, from which it is clear that the local SGE scores of the signaling genes abruptly increase in a collective manner around day 20. For MHCs-to-HCCs data, as shown in Fig. 3B, there is an obvious change in the network structure at embryonic day 12.5 (E12.5), signaling the cell fate transitions of the differentiation into hepatocytes and cholangiocytes after E12.5 [22]. The whole dynamics of signaling-gene network across all 7 time points is presented in Supplementary_material_1 (Figure S2A). Therefore, the network signature of a critical transition during embryonic cell differentiation is illustrated, which may benefit the understanding of molecular associations among cell populations. Moreover, to show the global view of the signaling genes, the landscape of local SGE scores was presented in Fig. 3E, in which the peak of local SGE scores for signaling genes appeared at E12.5. For hESCs-to-DECs data, there is a drastic change in the network structure at 36 h (Fig. 3C), signaling the cell fate transitions of the differentiation induction into the definite endoderm at 72 h [21]. The dynamical evolution of the PPI network across all 6 time points is provided in Supplementary_material_1 (Figure S2B). Moreover, to show the evolution of the signaling genes in a global view, the landscape of local SGE score is presented in Fig. 3F, in which the peak local SGE of signaling genes appears at 36 h. Clearly, by exploring the dynamical change of gene associations, SGE offers an insight of critical transition during the embryonic differentiation from the perspective of network dynamics.

**Figure 3:**
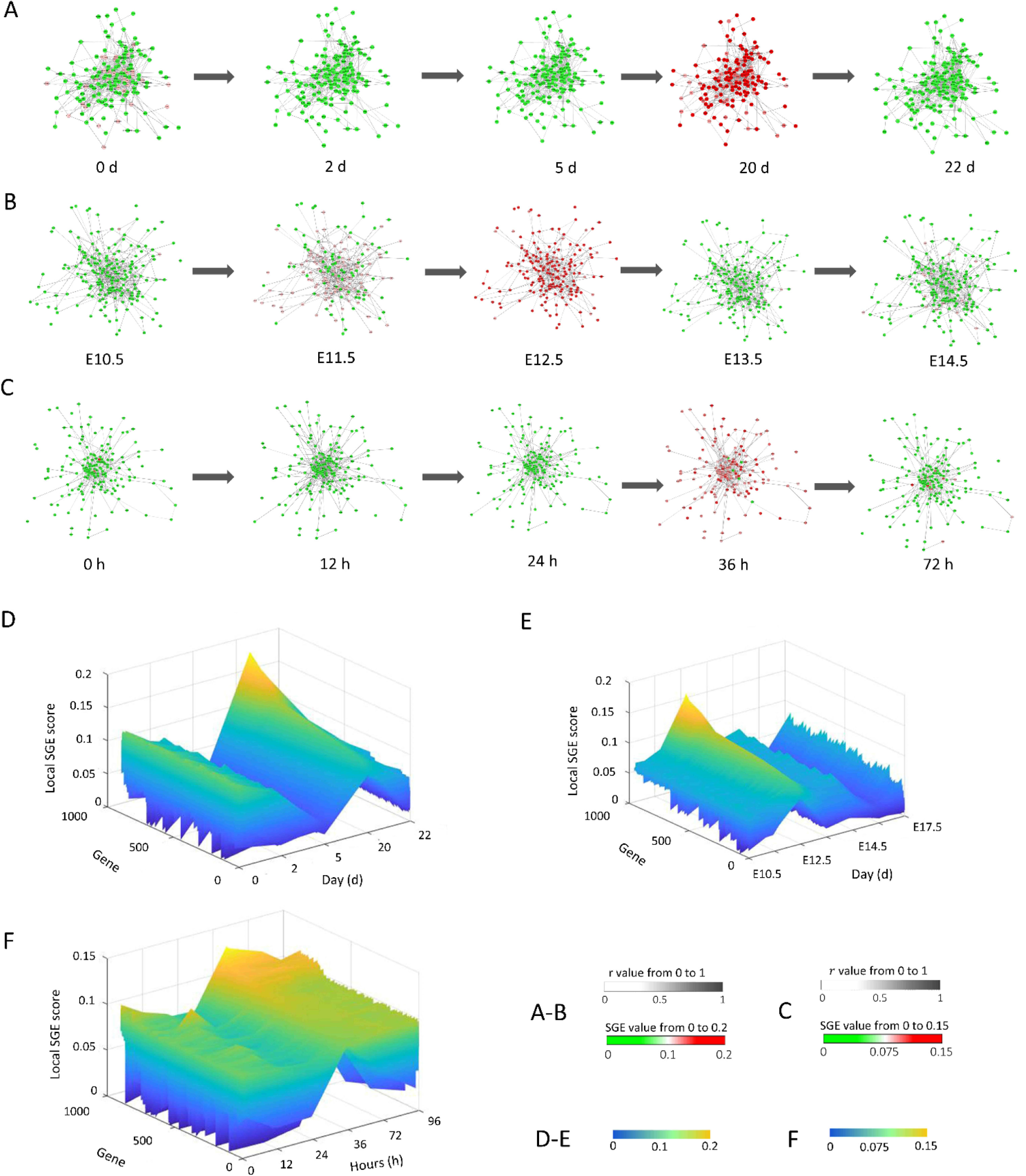
The dynamic evolution of gene regulatory networks and local SGE scores. Based on the SGE method, the key gene regulatory networks were reconstructed for the signaling genes (top 5% genes with the largest local SGE score) based on scRNA-seq data, where the color of each node represents the mean local SGE value (Eq.(2)) and the color of each edge represents the statistical dependency index (*r* in Eq.(1)). The dynamical evolution of gene regulatory networks for **(A)** MEF-to-Neurons data, which illustrates a significant change of the gene-gene associations at day 5 during the embryonic differentiation from MEF to neurons; **(B)** MHCs-to-HCCs data, in which the most significant change of the gene-gene associations occurs at E12.5. **(C)** hESCs-to-DECs data, which shows a significant change appearing at 36 h. The landscape of local SGE scores illustrates the dynamic evolution of network entropy in a global view for **(D)** MEF-to-Neurons data, **(E)** MHCs-to-HCCs data, and **(F)** hESCs-to-DECs data.

### 3.3 Temporal clustering and pseudo-trajectory analysis

The data transformation from the gene expression matrix to the SGE matrix not only helps to detect the critical transitions of embryonic development, but provides a better way to perform clustering analysis on cells during a biological process and thus explore dynamical information of cell populations. The t-distributed stochastic neighbor embedding (t-SNE), a nonlinear method to perform dimension-reduction [25], is applied to carried out dimension-reduction analysis and visualization, which has been extensively used in the analysis of scRNAseq data. A group of biomarkers are composed by top 5% genes with the largest local SGE value and top 5% genes with the smallest local SGE value in tipping point. We compare the clustering performance between SGE and gene expression (EXP) based on biomarkers. For MEF-to-Neuron, MHCs-to-HCCs and hESCs-to-DECs data, the clustering analyses are shown in Figs. 4A-4B, Figs. 4D-4E and Figs. 4G-4H, the clustering analysis based on SGE can distinguish the state of cells at different time points while the gene expression fails. Moreover, from the results as shown in the Figure S3 of Supplementary_material_1, the SGE method succeeded in distinguishing different cell types in three states, i.e., before-transition, critical-transition and after-transition state, but the gene expression fails to make such distinction. The result of dimension-reduction and visualization for NPCs-to-Neurons data and mESCs-to-MPs data is given in Supplementary_material_1 (Figure S4). Besides, the heatmap of SGE value for biomarkers stratified by three states (before-transition, critical-transition and after-transition state) while the heatmap of gene expression value fails (see Supplementary_material_2 for details). The best possible clustering analysis result of all datasets are obtained from the SGE method, which illustrates that SGE has a superior performance than the original gene expression.

**Figure 4:**
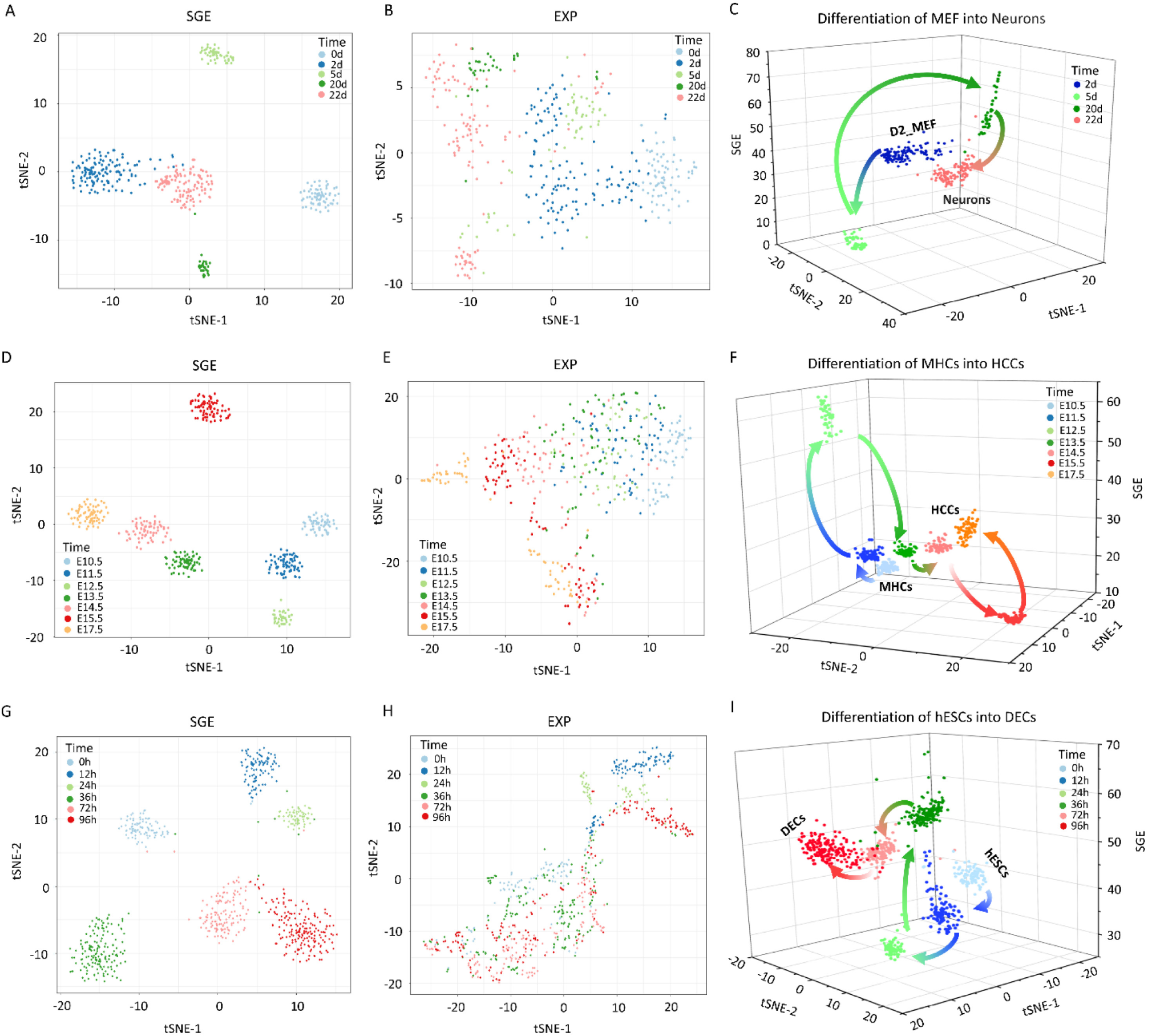
Comparison of clustering performance and pseudo-temporal trajectories of cell differentiation. Temporal clustering performance (t-SNE) between SGE and EXP and the differentiation trajectories for **(A)-(C)** MEF-to-Neurons data, **(D)-(F)** MHCs-to-HCCs data, and **(G)-(I)** hESCs-to-DECs data. Nodes in different colors represent cells from different time points. Clearly, SGE distinguishes the temporal cell state better than EXP. The differentiation trajectories can be accurately predicted by SGE scores.

To further validate the SGE performance, the pseudo-trajectory analysis was performed on the scRNA-seq data. Based on the temporal cell clustering by SGE, the three-dimensional representations of cell-lineage trajectories for three datasets are shown in Fig. 4C, Fig. 4F and Fig. 4I. The *z*-axis represents SGE potency estimation, while the *x* and *y* axes correspond to the t-SNE components. In Fig. 4C, we present the differentiation trajectories from MEF to neurons where MEF differentiated into neurons after 20 days. For MHCs-to-HCCs data, SGE predicted the dynamic differentiation trajectories from MHCs to HCCs (Fig. 4F). The MHCs-to-HCCs transition occurs immediately after embryonic day 12.5 (E12.5), which is consistent with the results of the original experimental observation [22]. Thus, the dynamics of cell fate decisions from MHCs to HCCs are revealed by such pseudo-temporal trajectories of SGE score, characterizing the underlying critical transition of the biological system during early embryonic development. When applied to hESCs-to-DECs data, the developmental trajectories of cell differentiation from hESCs to DECs are shown in Fig. 4I. The differentiation toward DECs appears after 36 h, which coheres with the experimental results [21]. These results demonstrate that the SGE-based potency estimation can track the dynamic changes in cell potency, as well as the specific time point at which the cell fate commitment or the differentiation into distinct cell types occurs.

### 3.4 Discovering “dark genes” by SGE score

In the biomedical field, differentially expressed genes play important roles in finding new biomarkers, key regulators and drug targets. However, some non-differentially expressed genes may also be involved in the essential biological processes, and should not be ignored. Actually, references showed that such genes are enriched in key functional pathways and performs well in prognosis [26] and may play biological roles in endothelial cells (EC) [12]. During the analysis of the above single-cell datasets, some genes were also discovered as the “dark” genes, which were non-differential in gene expression, but sensitive to SGE scores. These genes show a significant difference between the critical point and non-critical point at the network level, rather than at the gene expression level. We performed the differential SGE analysis on the five embryonic time-course differentiation datasets. The SGE and gene expression (EXP) were compared based on the signaling genes (top 5% genes with the largest local SGE score). Figures 5A-5C showed some “dark genes” of MEF-to-Neuron, MHCs-to-HCCs, and hESCs-to-DECs data. Other “dark genes” for these three datasets were respectively presented in Supplementary_material_3, Supplementary_material_4, and Supplementary_material_5. The results for the mESCs-to-MPs data and NPCs-to-Neurons data are respectively provided in Supplementary_material_6 and Supplementary_material_7. It is obvious that there are no significantly differential changes at the gene expression level, but significantly differential changes at the network entropy (SGE) level. Some “dark genes” have been reported to be associated with embryonic development, which illustrates that these “dark genes” play important roles in embryonic development. For these three datasets, the “dark genes” which are associated with embryonic development are demonstrated in Table 1-3, respectively.

**Table 1.** The information of important “dark genes” in MEF-to-Neurons data.

| Gene | Location | Family | Relation with embryonic development |
| --- | --- | --- | --- |
| **POLDIP2** | Cytoplasm | other | POLDIP2 knockout results in perinatal lethality, reduced cellular growth and increased autophagy of mouse embryonic fibroblasts [27]. |
| **HINT1** | Nucleus | enzyme | Deletion of the HINT1 gene enhances the growth of mouse embryonic fibroblasts [28]. |
| **EIF4G2** | Cytoplasm | translation regulator | EIF4G2 expression plays an important role in mouse embryo development [29] (Buim et al., 2005). |
| **EGLN1** | Cytoplasm | enzyme | EGLN1 is required for embryonic development [30]. |
| **DAD1** | Cytoplasm | other | Deletion of DAD1 in mouse embryo development induces an apoptosis-associated embryonic death [31]. |
| **BECN1** | Cytoplasm | other | The MiR-291a/b-5p can inhibit autophagy by targeting BECN1 during mouse preimplantation embryo development [32]. |
| **BAG1** | Cytoplasm | other | BAG1 is essential for differentiation and survival of hematopoietic and neuronal cells [33]. |
| **PPP2R2D** | Nucleus | other | PPP2R2D is correlated with embryonic growth and development [34]. |

**Table 2.**
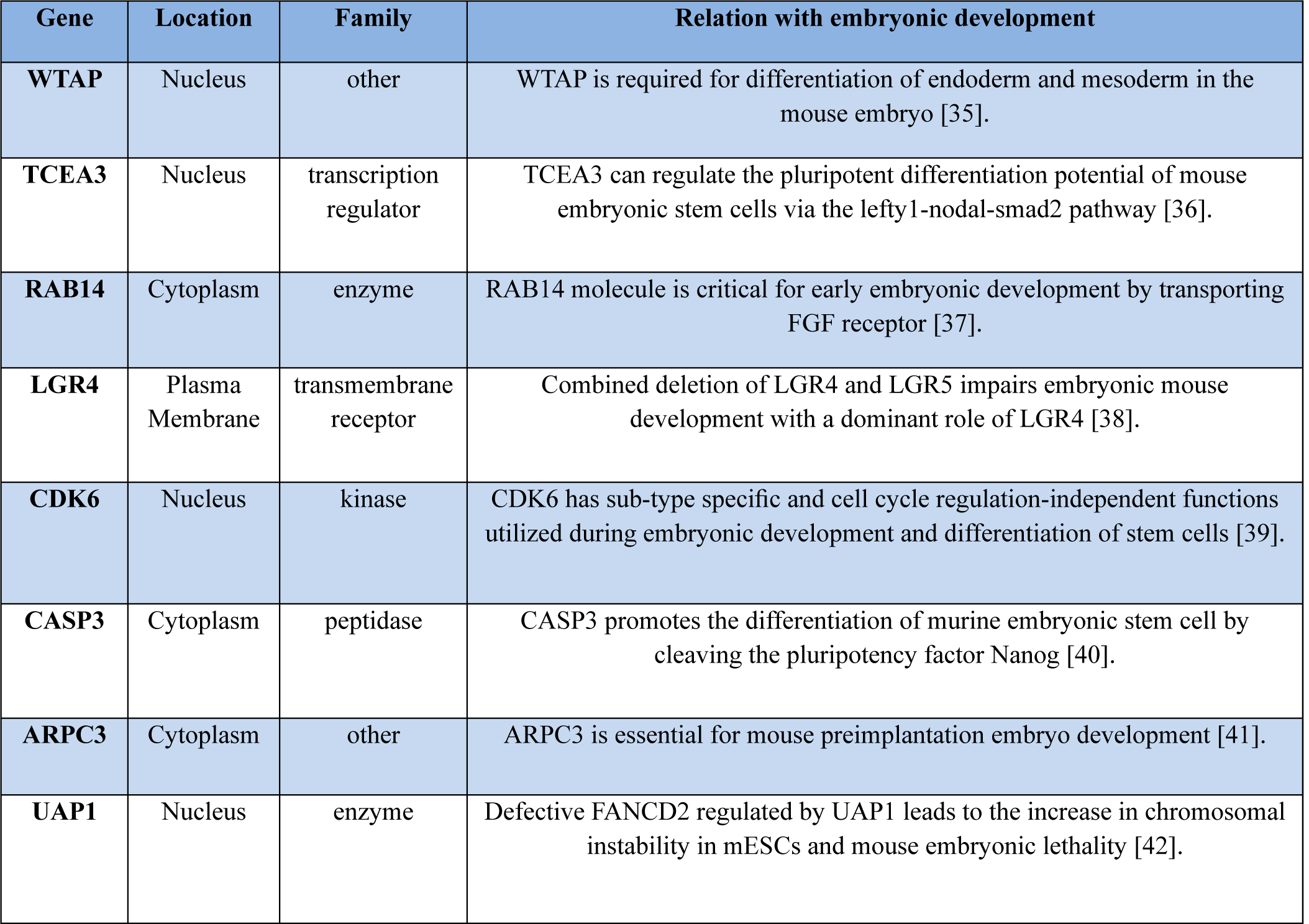
The information of important “dark genes” in MHCs-to-HCCs data.

**Table 3.** The information of important “dark genes” in hESCs-to-DECs data.

| Gene | Location | Family | Relation with embryonic development |
| --- | --- | --- | --- |
| **TMOD1** | Cytoplasm | enzyme | Absence of TMOD1 in differentiating embryonic stem cells leads to delayed myofibril assembly [43]. |
| **DOK4** | Plasma Membrane | other | DOK4 plays an important role during embryonic dorsal root ganglia neurons development [44]. |
| **ZEB2** | Nucleus | translation regulator | ZEB2 expression is essential for embryonic hematopoietic stem and progenitor cell (HSPC) differentiation in the fetal liver [45]. |
| **ZNF521** | Nucleus | transcription regulator | ZNF521 can efficiently drive embryonic stem cells to neural progenitors [46]. |
| **ZIC2** | Nucleus | transcription regulator | ZIC2 is an enhancer-binding factor required for embryonic stem cell specification [47]. |
| **VPS41** | Cytoplasm | transporter | VPS41 is essential for embryonic development [48]. |
| **SCML2** | Nucleus | transcription regulator | SCML2 plays an essential role in the modulation of self-renewal and differentiation of embryonic stem (ES) cells [49]. |
| **FGF2** | Extracellular Space | growth factor | FGF2 switches the outcome of BMP4-induced human embryonic stem cell differentiation [50]. |

**Figure 5:**
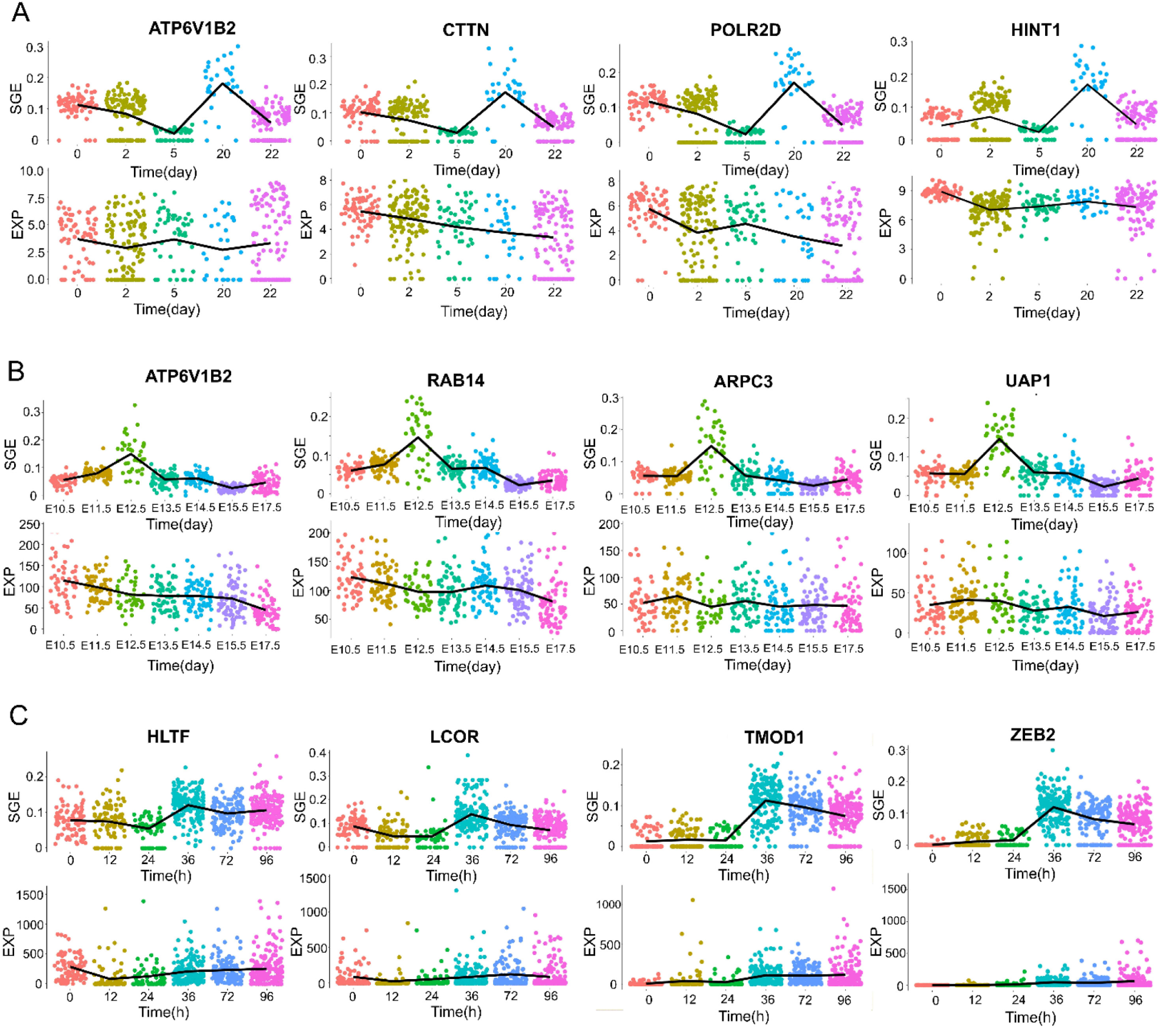
The embryonic time-course differentiation analysis based on “dark genes”. **(A)** POLR2D, ATP6V1B2 and CTTN, and HINT1. **(B)** WTAP, RAB14, ARPC3, and UAP1. **(C)** HLTF, LOCR, TMOD1, and ZEB2. It is obvious that there has no significantly differential changes at the gene expression level, but significantly differential changes at the SGE level. The SGE of dark genes show their peaks at the tipping point, which reveals embryonic development.

### 3.5 Revealing vital biological signals by common dark genes

Based on genes with differential SGE values, we found 6 common signaling genes (CSGs) for human embryo development among NPCs-to-Neurons data and hESCs-to-DECs data (Figure S6A of Supplementary_material_1) and other 14 among the mouse embryo development datasets (the Figure S6B of Supplementary_material_1). To evaluate their function in embryo development, the Reactome and KEGG pathway enrichment analysis is performed for these overlap genes.

For NPCs-to-Neurons data and hESCs-to-DECs data, it has been confirmed that common signaling genes, such as LOCR and HLTF (Fig. 5C), play a relatively important role in embryonic differentiation. LOCR, as an important molecule in the phosphatidylinositol signaling system, acts as a signal transduction element in consensus genes and may also participates in the regulation of TNFR1 signaling, interacts with the TNFR1-induced NFkappaB signaling pathway, and activates tumor necrosis factor receptor 1 (TNFR1). Multiple signal transduction pathways can be triggered to induce inflammation, cell proliferation, survival, or cell death [51-53]. At the same time, to respond to a wide range of extracellular stimuli, thereby promoting differentiation, proliferation, cell motility, cell survival, and some other important cellular behavior [54-56], LOCR and HLTF together act as the RAF / MAP kinase cascade element in the RAS-RAF-MEK-ERK pathway to participate in controlling downstream MAPK1 / MAPK3 signaling by directly activating MAP2K and MAPK, and MAPK3 and MAPK1 will be phosphorylated by MAP2Ks 1 and 2.

In addition, LCOR participates in TCF dependent signaling in response to WNT signal together with MGA. The WNT pathway is one of the most important signaling pathways in cells for cell proliferation. In the classical WNT signaling pathway, the binding of WNT ligands to frizzled protein (FZD) and lipoprotein receptor-related protein (LRP) receptors leads to the destruction of complex inactivation, the stabilization and nuclear translocation of β-catenin and subsequent activation of TCF/LEF-dependent transcription. Transcriptional activation in response to classical WNT signaling controls cell fate, stem cell proliferation, and self-renewal, and promotes tumorigenesis [57-59].

As an important transcription factor, HLTF has both helicase and E3 ubiquitin ligase activities. We have noticed that it is directly involved in Ras activation upon Ca2 + influx through the NMDA receptor [60]. Ras catalyzes its effector substrate to regulate a series of important functions related to cell growth, differentiation, and apoptosis. Besides, HLTF, together with MAG, also plays an important role in the cell cycle. Also, as described in the GSE102066 article [20], HLTF is also directly involved in the neurobiological process of negative regulation of NMDA receptor-mediated neuronal transmission, which might also be one of the key regulators of brain / spinal neuron differentiation after 24 hours. It should be noted that the role of these gene products in the pathway also belongs to the upstream of signaling. For example, that LOCR and HLTF play a direct role in controlling downstream MAPK pathway when they participate the RAF / MAP kinase cascade signal cascade process. At the same time, this kinase cascade, as a downstream effector of FLT3 Signaling, communicates FLT3 Signaling with the MAPK pathway. Beyond that, RAF / MAP kinase cascade is also important in CREB1 phosphorylation through NMDA receptor-mediated activation of RAS signaling, which may also lead cell proliferation.

Among the 14 common signaling genes across MHCs-to-HCCs, MEF-to-Neurons and mESCs-to-MPs datasets, it has been seen that some genes, including POLR2D, ATP6V1B2 and CTTN (Figs. 5A-5B), also participate in mouse embryonic differentiation. POLR2D directly participates in RNA Polymerase II transcription initiation as the main component of RNA polymerase 2, which is a necessary step for gene expression. The formation of an open complex exposes the template strand to the catalytic center of RNA polymerase II. This will promote the formation of the first phosphodiester bond, which marks the start of transcription [61]. The initiation of transcription is the main regulatory point of gene expression [62]. As well-known already, in the absence of the transcription process, the development of early embryonic cells generally depends on the mRNA inherited mother [63]. This means that the initiation of transcription may indicate that the embryonic cell initiates their own distinct transcription process, and the embryonic cell officially enters the autologous development process. This is exactly consistent with the pluripotent withdrawal process mentioned in the GSE79578 article [22], so POLR2D may contribute to this process. Besides, ATP6V1B2 directly participates in the amino acid-activated mTOR receptor pathway by participating in processing upstream amino acid stimulation signals and transmitting to the regulator and then activating the downstream mTOR effector pathway. The mTOR can regulate neuronal proliferation, survival, growth, and function, this is crucial for the developmental process, and relaxing the control of mTOR at any stage of development may have harmful consequences [64]. This process may indicate that ATP6V1B2 may be a key gene for cell fate determination explored in the data article of GSE67310 [19]. At the same time, the CTTN gene is a part of the cell tight junction component, responding extracellular pressure and activating downstream actin assembly. The actin assembly dynamics is strictly controlled by time and space [65], while the actin-assembled cytoskeleton has various physiological and pathological functions for cell migration, differentiation, embryonic development [66]. Therefore, CTTN may play an important role in embryonic development by regulating actin assembly.

## 4. Discussion and conclusion

Predicting a cell fate or lineage transition for cell differentiation is a task of biological and clinical importance [11]. Understanding of such cell fate commitment may help to construct individual-specific disease modeling, and design therapies with great specificity for complex diseases relevant to cell differentiation [77]. Most of the existing methods applied in analyzing scRNA-Seq data are based on the gene expression and its statistical quantities. However, gene expressions are generally considered too unstable to characterize the dynamics of biological process [12,78,79]. In this study, we developed the SGE method to explore the dynamic information of gene-gene associations from scRNA-Seq data, and thus predict the cell fate transition during early embryonic development. The proposed method has been applied to five single-cell RNA sequencing datasets and successfully identified the critical stage or tipping point of the impending cell fate transition. For instance, the significant change of SGE score indicates the critical point (day 20) of MEF-to-Neuron data before the differentiated into induced neurons, the critical point (36 h) of hESCs-to-DECs data prior to the differentiation induction into definite endoderm (DE), and the tipping point (E12.5) of MHCs-to-HCCs data before the differentiation into hepatocytes and cholangiocytes.

By transforming the sparse gene expression matrix from the scRNA-seq data into a non-sparse graph entropy matrix, SGE offers a new computational insight to the single cell analysis, and helps to discover the signal of the underlying cell fata commitment. Firstly, for each gene, SGE method provides a gene-specific local SGE score, which transforms the data from unstable gene expression form to relatively stable network entropy (SGE) form. Therefore, rather than the originally measured gene expression data, we use the transformed SGE for further analysis, which can reliably characterize the status of the dynamical biological process. The analysis results in this study illustrate the better performance of SGE than the original gene expression in both indicating critical transitions and cell clustering of temporal information. Secondly, the change of SGE scores also identifies the pseudo-temporal trajectories of cell differentiation, which helps to analyze the differentiation potency of cells. Clearly, the dynamics of cell fate decisions are revealed by such SGE-based trajectories, thus characterizing the underlying critical transition of the biological system during early embryonic development. Besides, SGE helps to uncover “dark genes”, which are non-differential in gene expression but sensitive to SGE score. Such non-differential genes were often ignored by the traditional differential gene expression analyses. However, some non-differential genes may also be involved in the key biological activities of cells and play important roles in embryonic development. Notably, the SGE method is model-free, that is, the SGE strategy requires neither feature selection nor model/parameter training. In summary, SGE opens a new way to predict a cell fate transition at the single-cell level, which is helpful in tracking cell heterogeneity and elucidating molecular mechanism of embryonic cell differentiation.

## List of abbreviations

scRNA-Seq: single-cell RNA sequencing:
SGE: single-cell graph entropy:
EXP: the gene expression:
MEF: mouse embryonic fibroblasts:
HCCs: hepatocytes and cholangiocytes cells:
NPCs: neural progenitor cells:
hESCs: human embryonic stem cells:
DECs: definitive endoderm cells:
MHCs: mouse hepatoblasts cells:
mESCs: mouse embryonic stem cells:
DE: definite endoderm:
MPs: mesoderm progenitors:
DNB: dynamic network biomarker:
iN: induced neuron:
EC: endothelial cells:
ES: embryonic stem:
t-SNE: t-distributed stochastic neighbor embedding:
HSPC: hematopoietic stem and progenitor cell:
CSGs: common signaling genes:
DEGs: differential expressed genes:

## Acknowledgments

This work was supported by National Natural Science Foundation of China (Nos. 11771152, 11901203, 11971176), Guangdong Basic and Applied Basic Research Foundation (2019B151502062), China Postdoctoral Science Foundation funded project (No. 2019M662895) and the Fundamental Research Funds for the Central Universities (2019MS111).

## Author Contributions Statement

R.L. and P.C conceived the project; J.Z, C.H., and X.Z. performed computational and analysis. All authors wrote the manuscript. All authors read and approved the final manuscript.

